# Computational Design and Analysis of Modular Cells for Large Libraries of Exchangeable Product Synthesis Modules

**DOI:** 10.1101/2021.03.15.435526

**Authors:** Sergio Garcia, Cong T. Trinh

## Abstract

Microbial metabolism can be harnessed to produce a large library of useful chemicals from renewable resources such as plant biomass. However, it is laborious and expensive to create microbial biocatalysts to produce each new product. To tackle this challenge, we have recently developed modular cell (ModCell) design principles that enable rapid generation of production strains by assembling a modular (chassis) cell with exchangeable production modules to achieve overproduction of target molecules. Previous computational ModCell design methods are limited to analyze small libraries of around 20 products. In this study, we developed a new computational method, named ModCell-HPC, capable of designing modular cells for large libraries with hundredths of products with a highly-parallel and multi-objective evolutionary algorithm. We demonstrated ModCell-HPC to design *Escherichia coli* modular cells towards a library of 161 endogenous production modules. From these simulations, we identified *E. coli* modular cells with few genetic manipulations that can produce dozens of molecules in a growth-coupled manner under different carbons sources. These designs revealed key genetic manipulations at the chassis and module levels to accomplish versatile modular cells. Furthermore, we used ModCell-HPC to identify design features that allow an existing modular cell to be re-purposed towards production of new molecules. Overall, ModCell-HPC is a useful tool towards more efficient and generalizable design of modular cells to help reduce research and development cost in biocatalysis.

## 1 Introduction

Modular design has gained recent interest as an effective approach to understand and redesign cellular systems.^1^ In the fields of metabolic engineering and synthetic biology, various modularization strategies^2–7^ have been proposed to address the slow and expensive design-build-test cycles of developing microbial catalysts for renewable chemical synthesis.^8^ A promising system-level modularization^9^ approach is ModCell,^4^ that aims to design a modular (chassis) cell compatible with exchangeable production modules that enable metabolite overproduction. ModCell could be used as an effective tool to design modular cells capable of efficiently producing a vast number of molecules offered by nature with minimal strain optimization requirements,^10,11^ but it remains unexplored for large product libraries.

Previous efforts in computational modular cell design are limited to analyze small libraries of around 20 products.^4,6^ However, the design of modular cells for larger product libraries is both of practical and theoretical interest. Theoretically, using large libraries can lead to more general modular cell design rules, which might help to explain the naturally existing modularity of metabolic networks.^1^ Practically, such modular cells could be implemented with genetic engineering techniques that enable rapid pathway generation, such as combinatorial ester pathways,^12^ and where the modular cell could serve as a versatile platform for pathway selection and optimization using adaptive laboratory evolution.^13^

Modular cell design was formulated as a multi-objective optimization problem (MOP), named ModCell2, where each target phenotype activated by a module is an independent objective.^4^ ModCell2 was solved with multi-objective evolutionary algorithms (MOEAs) that used a master-slave parallelization scheme, where the objective functions are evaluated in parallel by slave processes, but every other step in the algorithm is performed serially (Figure 1 a).^4,5^ This approach contains many serial steps, and hence limits the scalability of the algorithm with the number processes according to Ahmdal’s law.^14^ In particular, large population sizes, an effective strategy to deal with many objectives,^5,15^ can dramatically slow down serial algorithm operations such as non-dominated sorting in NSGA-II,^16^ one of the best performing MOEAs to solve ModCell2.^5^ Furthermore, increasing the product library size for ModCell leads to very large multi-objective optimization problems, which are notoriously difficult to solve.^17,18^ Therefore, the master-slave approach used in ModCell2 is not suitable to analyze large problems that contain hundredths of exchangeable production modules. A new parallelization approach that uses high-performance computing (HPC) more effectively is needed to advance ModCell.

**Figure 1:**
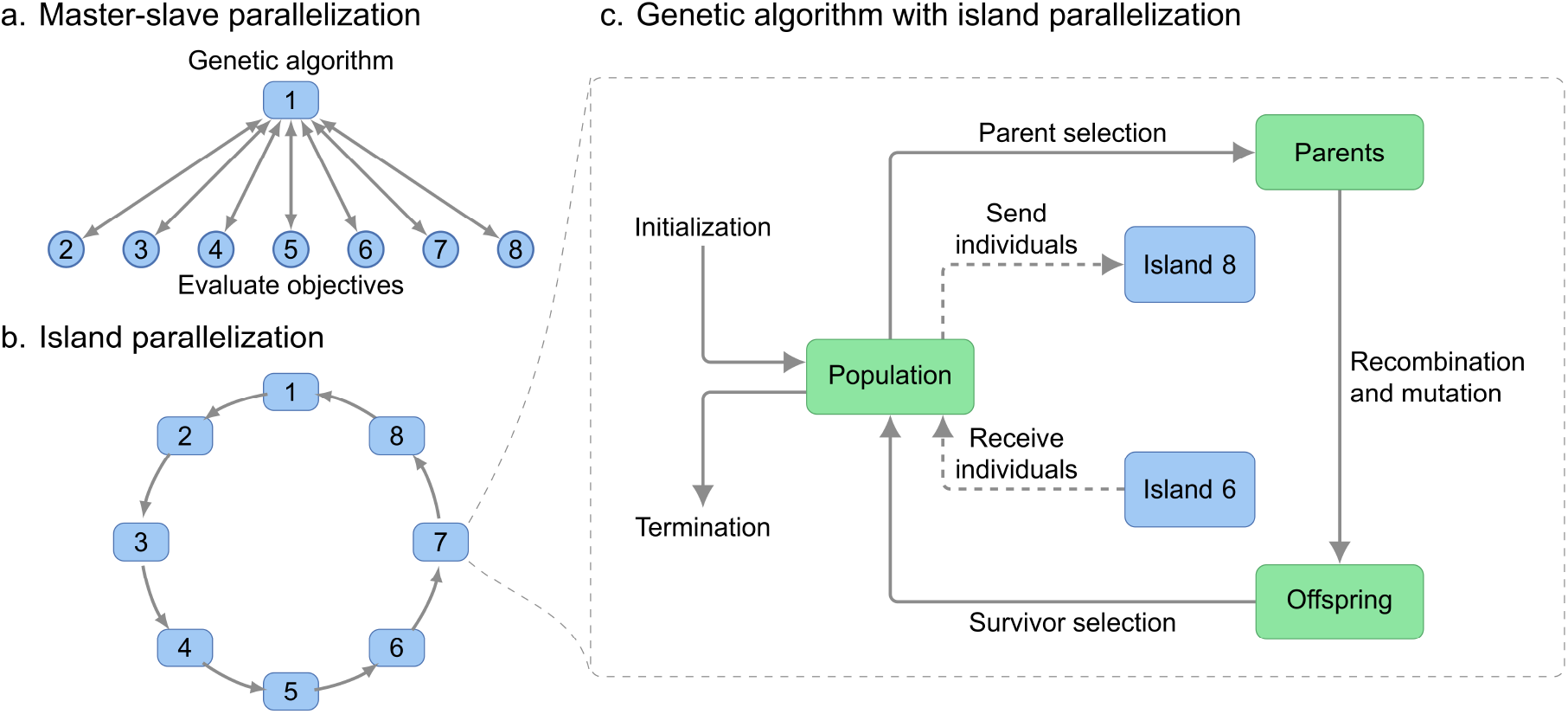
Parallelization schemes for multi-objective evolutionary algorithms. (a) Master-slave approach used in the original ModCell2 implementation. (b) Island parallelization following ring topology implemented in ModCell-HPC. (c) Key steps in the evolutionary algorithm.

In recent years, multiple approaches to harness HPC have been developed to solve single-objective evolutionary algorithms (EA).^19^ In particular, the island-parallelization approach has been proposed, where multiple instances of the EA are run independently but communicate with each other to enhance overall convergence towards optimal solutions (Figure 1 b). This new approach helps address the serial bottlenecks of the master-slave approach by separating the algorithm into highly independent processes that directly map to the computing hardware. While this approach has not been throughly examined in MOEA, there are a few successful applications to specific design problems.^20–22^

In this study, we developed ModCell-HPC, a highly parallel MOEA that uses the island parallelization approach to solve modular cell design problems with hundredths of objectives. We demonstrated ModCell-HPC to design *Escherichia coli* modular cells with a large production module library of metabolically and biochemically diverse endogenous compounds. Analysis of these designs revealed key genetic manipulations both at the chassis and module levels required for highly compatible modular cells. Furthermore, we designed modular cells for conversion of various hexoses and pentoses, since these sugars are the main components of biomass feedstocks.^23^ Finally, we used ModCell-HPC to identify the features of a modular cell that makes it compatible towards new production modules.

## 2 Methods

### 2.1 Multi-objective optimization formulation of modular cell design problem

The modular (chassis) cell is built in a top-down manner by removing metabolic functions from a parent strain, and then inserting exchangeable modules into the chassis to create production strains that optimally display the target phenotypes. Due to the conflicting biochemical and metabolic requirements of different product synthesis pathways, the modular cell design problem is formulated as the following MOP known as ModCell2:^4^

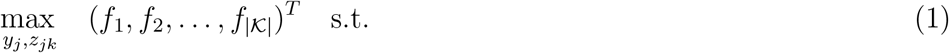

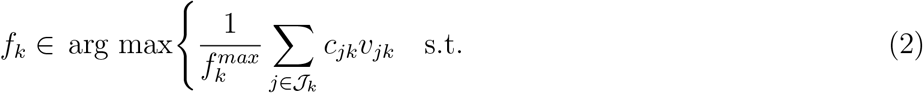

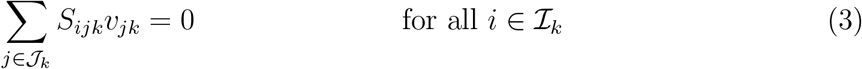

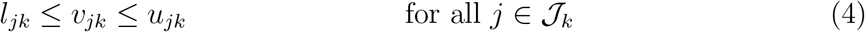

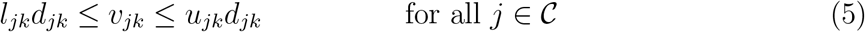

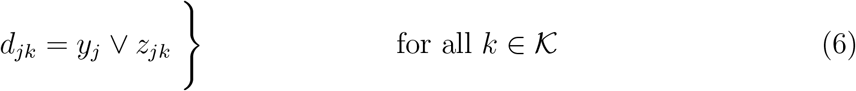

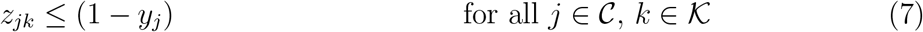

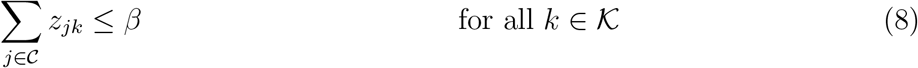

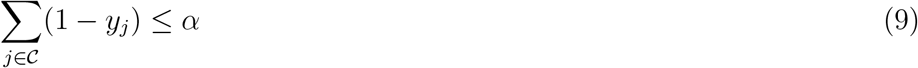

This MOP simultaneously maximizes all objectives *f_k_* (1), where *k* belongs to the set of production networks 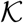. Each production network represents the combination of the chassis with a specific production module, and it is simulated through a stoichiometric model^24^ (2–6) with a set of metabolites 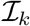 and a set of reactions 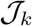. The stoichiometric model predicts metabolic fluxes according to the following constraints: (i) mass-balance (3), where *S_ijk_* represents the stoichiometric coefficient of metabolite *i* in reaction *j* of production network *k*, (ii) flux bounds (4) that determine reaction reversibility and available substrates, where *l_jk_* and *u_jk_* are lower and upper bounds respectively, and (iii) genetic manipulation (5), i.e., deletion of a reaction *j* in the chassis through the binary indicator *y_j_*, or insertion of a reaction *j* in a specific production network *k* through the binary indicator *z_jk_*. Only a subset of all metabolic reactions, 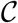, are considered as candidates for deletion, since many of the reactions in the metabolic model cannot be manipulated to enhance the target phenotype.

The desirable phenotype *f_k_* for production module *k* is determined based on key metabolic fluxes *υ_jk_* (mmol/gDCW/h) predicted by the model (2–5). For this study, we selected the weak growth coupled to product formation (*wGCP*) design objective that requires a high minimum product synthesis rate at the maximum growth rate, enabling growth selection of optimal production strains. Hence, in *wGCP* design, the inner optimization problem seeks to maximize growth rate while calculating the minimum product synthesis rate through the linear objective function (2). Here *Cj_k_* is 1 and –0.0001 for *j* corresponding to the biomass and product reactions across all networks *k*, respectively, and 0 otherwise. In general, the definition of *f_k_* needs not be linear and other design phenotypes can be defined.^4^

Finally, design constraints (7–9) define the limitations of the design variables representing genetic manipulations, *y_j_* and *z_jk_*. As part of modular cell design, reactions can be removed from the chassis but inserted back to specific production modules, enabling the chassis to be compatible with a broader number of modules (7). The total numbers ofl module reaction additions and reaction deletions in the chassis are limited by parameters *β* (8) and *α* (9), respectively.

To define the solutions of ModCell2 (1–9), the general multi-objective optimization problem with design variables *x* from a set ***χ*** and objective functions *f_i_*(*x*) is expressed as follows:

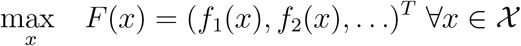

The solution of such an optimization problem is denoted as a Pareto set:

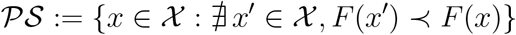

Here *F*(*x′*) ≺ *F*(*x*) indicates that the objective vector *F*(*x′*) *dominates F*(*x*), defined as *f_i_*(*x′*) ≥ *f_i_*(*x*) for all objectives *i*, and *f_i_*(*x′*) ≠ *f_i_*(*x*) for at least one *i*. Hence, the Pareto set contains all non-dominated solutions to the optimization problem; that is, when comparing any two non-dominated solutions, the value of a certain objective must be diminished in order to increase the value of a different objective. The projection of the Pareto set on the objective space is denoted as a Pareto front:

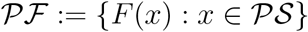

### 2.2 Implementation of many-objective evolutionary algorithm with high-performance computing

To overcome the issues of the master-slave approach (Figure 1 a) used in ModCell2,^4^ we implemented an island parallelization scheme,^19^ where each computing process is an instance of the MOEA (Figure 1 b). These instances exchange individuals (i.e., potential solutions) in a process called migration, hence enhancing overall convergence towards optimal solutions (Figure 1 c). The migration operation can be performed in different modes, depending on which individuals from the local population are exchanged, and also how often such exchanges happen. These options are captured by the migration type and migration interval parameters, respectively (Table 1). To enhance performance and scalability, the migration process was implemented asynchronously, i.e., the population within each island can continue to evolve without a need to wait for sent individuals to arrive at their destination island or for incoming individuals to be received.

**Table 1:** Island-MOEA parameters evaluated in ModCell-HPC.

| Name | Description |
| --- | --- |
| Population size | Number of individuals per island. |
| Migration type | “ReplaceBottom”: After non-dominated sorting of the Pareto front<sup>16</sup> (survivor selection), top individuals are sent and bottom individuals replaced. “Random”: Random individuals are sent and replaced. |
| Migration interval | Number of generations between migration events. |
| Run time | Wall-clock time for which the MOEA runs. It determines the total number of generations. |
| Cores | Each island is a computing core at the hardware level. |

To improve the quality of the MOEA solutions, we implemented two post-processing steps specific to ModCell (Figure S1): First, we eliminate *futile module reactions*. These module reactions once removed do not diminish the objective value of the associated production network. Second, we coalesce multiple designs with the same deletions but different module reactions. This combination helps obtain a superior solution.

The software implementation of the proposed island-MOEA, denoted *ModCell-HPC*, is written in the C programming language and available at https://github.com/TrinhLab/modcell-hpc.

### 2.3 Computation hardware

We conducted all ModCell-HPC computations in *beacon* nodes from the Advanced Computing Facility at the Joint Institute for Computational Science, The University of Tennessee and Oak Ridge National Laboratory. Each node contains a 16 core Intel Xeon E5-2670 central processing unit (CPU) and 256 GB of random access memory (RAM). The results were analyzed in a desktop computer with an Intel Core i7-3770 CPU and 32 GB of RAM.

### 2.4 Target product identification

The target products are endogenous *E. coli* metabolites that meet the following requirements: i) their maximum theoretical yields are above 0.1 (mol product/mol of substrate); ii) they are organic; and iii) they could be produced anaerobically in a growth coupled manner with a yield above 50%, a property determined in a previous study.^25^ If a given metabolite meets all these conditions but appears in multiple compartments, only one location is choosen. Implementation of these selection criteria resulted in 161 target metabolites. Metabolites that did not have a secretion mechanism originally present in the model required an exchange pseudo-reaction that represents metabolite secretion to the growth medium or intracellular accumulation at steady-state. The products in the selected library have diverse molecular weights and are overall highly reduced (Figure S2).

### 2.5 Model configuration

We used the iML1515 *E. coli* model^26^ for all simulations. To configure the model, glucose uptake was set to 15 (mmol/gCDW/h); the default ATP maintenance value in iML1515 was used; 20% of the maximum anaerobic growth rate was used as the minimum growth rate, corresponding to 0.0532 (1/h); and only commonly observed fermentation products were allowed for secretion. This model configuration is equivalent to the previous modular cell design studies^4^ except for the higher glucose uptake rate. This rate was increased to match the study of Kamp and Klamt^25^ which was partially used here to identify target products.

### 2.6 Design characterization

#### 2.6.1 Compatibility

An important qualitative feature of a designed modular (chassis) cell is module compatibility. The chassis is *compatible* with a module if the performance of the resulting production strain is above a defined threshold of design objective value. In this study, we used the *wGCP* design objective that corresponds to the minimum product yield at the maximum growth rate,^4^ and selected a threshold of 0.5 to establish compatibility. Under these conditions, we expect a module compatible with the chassis can lead to a product yield above 50% of the theoretical maximum during the growth phase. The *compatibility* of a modular cell is defined as the number of modules that are compatible with it.

#### 2.6.2 Minimal covers

A *minimal cover* is the smallest group of modular cells needed to ensure all potentially compatible products in a library are compatible with at least one of the modular cells. To identify minimal (set) covers computationally, we use the classical integer programming formulation:

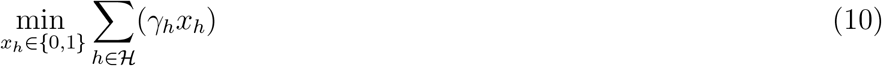

subject to:

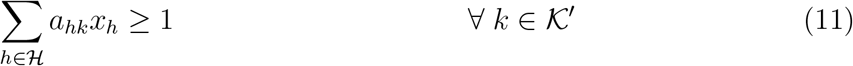

This optimization problem minimizes the number of designs in the set cover, where 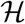 is the set of strain designs, *h*, produced by ModCell-HPC (10). The binary indicator variable *x_h_* takes a value of 1 if design *h* is selected as part of the set cover and 0 otherwise. Certain designs can be prioritized (e.g., they contain preferable genetic manipulations) using the weighting parameter γ*_h_*. However, we set γ*_h_* = 1 in all our simulations. All compatible products *k* must be included in at least one of the selected designs (11). The parameter *a_hk_* takes a value of 1 if product *k* is compatible with design *h* and 0 otherwise. There must exist at least one 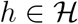 for which *a_hk_* = 1 to ensure a feasible solution exists; therefore, 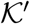 is the subset of products compatible in at least one design of 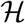.

To enumerate all minimal covers, we iteratively solved the minimal cover problem (10–11) with the addition, in each iteration, of an integer cut inequality (12) that removes a previously found solution 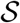.

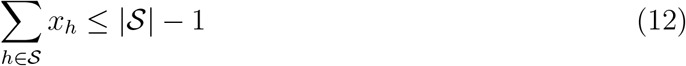

#### 2.7 Coverage performance indicator

Algorithm performance is tested against several parameter configurations, each producing a

Pareto front approximation 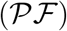. All resulting Pareto fronts are gathered into a reference

Pareto front 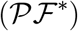. Coverage, *C*, is defined as the fraction of solutions in 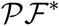 captured by a given approximation 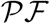:

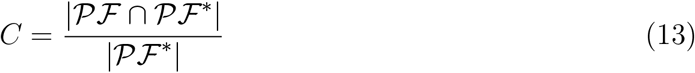

In our analysis, we only used unique non-dominated points in both 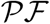 and 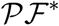 to avoid many alternative solutions from biasing the coverage indicator.

### 3 Results

#### 3.1 Tuning of ModCell-HPC method parameters

A known challenge of heuristic optimization approaches is their reliance on parameter tuning for rapid convergence towards optimal solutions. To identify sensible default parameters for ModCell-HPC, we first scanned parameter combinations with a previous 20-objectives problem^4^ that is fast to solve, then focused on the most relevant parameters for a large-scale problem with 161 objectives corresponding to the current product library. In both cases, we used two performance metrics to identify the best algorithm parameters: i) *Coverage*, that indicates the fraction of Pareto optimal solutions identified by a given parameter configuration (Section 2.7). ii) *minimal cover size*, i.e., the smallest number of modular cells needed to ensure all compatible products in the library that are compatible in at least one (Section 2.6.2). Coverage is a general and unbiased quantitative measure which is preferred over other similar metrics based on a previous study,^5^ while minimal cover size is based on practical goals.

In our initial benchmark study with the 20-objectives problem, we screened different total run times, migration interval, migration types, and population sizes (Table 1) for best achieving modular cell designs. The design parameters were set to *α* = 6 and *β* =1, which are sufficient to find highly compatible designs.^6^ For 1-hour run time, we observed the smallest population size (100) reached more generations (Figure S3 e,f) and hence achieved better results in both metrics (Figure S3 a,b). However, for a 2-hours run time, both population sizes of 100 and 500 attained similar cover sizes (Figure S3 g), indicating that a minimum of approximately 150 generations (Figure S3 e,f,k,l) is necessary for convergence of this problem, irrespective of the population size. Taken together, the different performance between 100 and 500 population sizes in relation to run time indicates that under limited run times an optimal population size could be found to attain sufficient generations for convergence. The migration interval only appeared detrimental at the highest value of 50 with the smallest population size of 100 at 1 hour (Figure S3 a,b,g,h); otherwise this parameter was considered secondary, and hence an intermediate value of 25 was selected for further simulations. Similarly, migration policy also appeared to be a secondary parameter; nonetheless, the “ReplaceBottom” migration policy was selected for further simulations since it is better or equal to the “Random” policy in all cases (Figure S3 c,d,i,j).

For the large-scale benchmark with 161 products, we investigated the importance of run time, population size, and the number of computational cores (Table 1). For this benchmark, the design parameters were set to *α* = 10 and *β* = 2 to enable successful designs without a large number of genetic modifications that can lead to unrealistic model predictions and implementation requirements. We evaluated 5 and 10 hour run times. For 5-hours run time, a population size of 200 was better in all metrics (Figure 2 a,b,c,e,f,g) and reached 50-100 generations (Figure 2 d). For a 10-hours run time, the population sizes of 200 and 300 had equivalent performance (Figure 2 e-g), despite the population size of 200 reaching approximately 50 generations more than the 300 population size. The population size of 100 underperformed at both run-times (Figure 2 a,b,e,f). Taken together, this large-scale benchmark study indicates that after a given number of generations, larger population sizes are comparable as long as they are above a minimum size. Hence, a population size of 200 is the minimum required for proper convergence and should be used under limited run times. Increasing the number of cores leads to more solutions (Figure 2 c,g), due to a larger meta-population (the total population of all islands). However, additional cores do not necessary find better solutions in terms of minimal cover size and individual product compatibility (Figure 2 b,f). These indicators plateaued at around 48 cores in both cases so this value was used for further simulations. Alternative communication topologies among islands^27^ may provide better scaling with cores but were not explored here.

**Figure 2:**
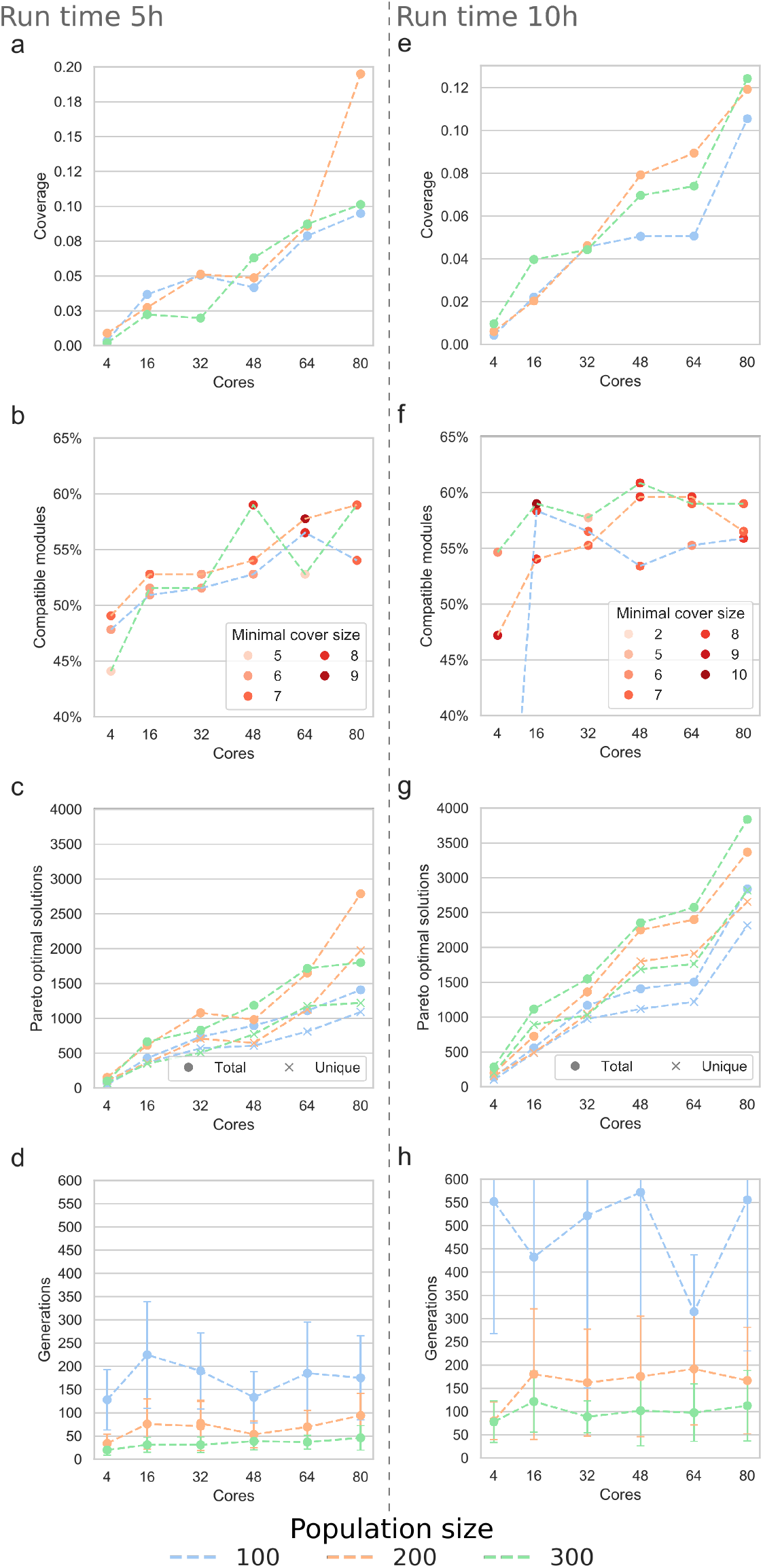
ModCell-HPC benchmark with 161 products. (a) and (e) Coverage is the fraction of Pareto optimal designs captured by a Pareto front approximation (Section 2.7). (b) and (f) Compatible modules indicates the products that appear in at least one design with a design objective above the compatibility threshold, while minimal cover size is the smallest number of designs needed to capture all compatible products (Section 2.6). (c) and (g) Total and unique number of solutions in the Pareto front approximations. (d) and (h) Total number of generations.

In summary, the benchmark performed here aims to provide a general guideline to use the ModCell-HPC. Furthermore, this parameter meta-optimization procedure can be repeated to fine-tune the algorithm to specific problem features (e.g., number of objectives) and computational resources available (e.g., run time and computing cores).

#### 3.2 Design of *E. coli* modular cells for large product library

##### A small number of genetic manipulations are sufficient for highly compatible modular cell

After tuning ModCell-HPC, we used it to design *E. coli* modular cells for our library of 161 products. First, we scanned a broad range of design parameter combinations (*α-β*: 5-1, 10-2, 20-4, and 40-8) to identify the required genetic manipulations for highly compatible designs (Figure S4 a). Increasing the number of genetic manipulations led to an average increase in design compatibility. However, the maximum compatibility remained around 50% of the library (80 products) for all cases. This result indicates that highly compatible modular cells can be built with a small number of genetic manipulations. We selected the designs with *α* = 5, *β* = 1 (Supplementary Material 2) for further analysis, since designs with few genetic manipulations are more accurately simulated and also better to implement in practice.

##### A few reaction deletions in central metabolism targeting byproducts and branch-points are key to build modular cells

We sorted reaction deletions according to how often they appear across designs (Table 2). The top 7 reactions are used ≥10% of the designs and belong to central metabolism, indicating their importance to accomplish growth-coupled-to-product-formation phenotypes. Overall, the role of these deletions can be classified into two functions: i) to eliminate major byproducts and ii) to alter key branch-points in metabolism that influence the pools of precursor metabolites, including carbon, redox, and energy precursors. The first type of manipulations is generally intuitive and often used in metabolic engineering strategies.^28^ The second type of manipulations are not commonly identified unless metabolic model simulations are used,^29–31^ even though the importance of targeting metabolic branch-points was noted early.^32^ An example of this second type observed in our designs is TPI deletion, that activates the methylglyoxal bypass,^33^ reducing the overall ATP yield resulting from glucose conversion into pyruvate. Lower ATP yield limits biomass formation hence redirecting carbon flow towards products of interest. While such strategies are not common, TPI deletion predicted by model simulations was successfully used for enhanced 3-hydroxypropionic acid production,^29^ and ATP wasting has recently been proposed to enhance production of certain molecules. ^34^ Another example of branch-point manipulation is PPC deletion, that has been shown to lower flux from lower glycolysis towards the TCA cycle,^35,36^ resulting in lower succinate production, and an increased pool of *pep*, pyruvate and acetyl-CoA. Additionally, PPC deletion to increase the *nadph* pool for production of flavonoids was predicted by model simulation and experimentally validated.^31^ In summary, design of highly compatible modular cells requires not only major byproduct removal, but also manipulation of key branch points in central metabolism.

**Table 2:**
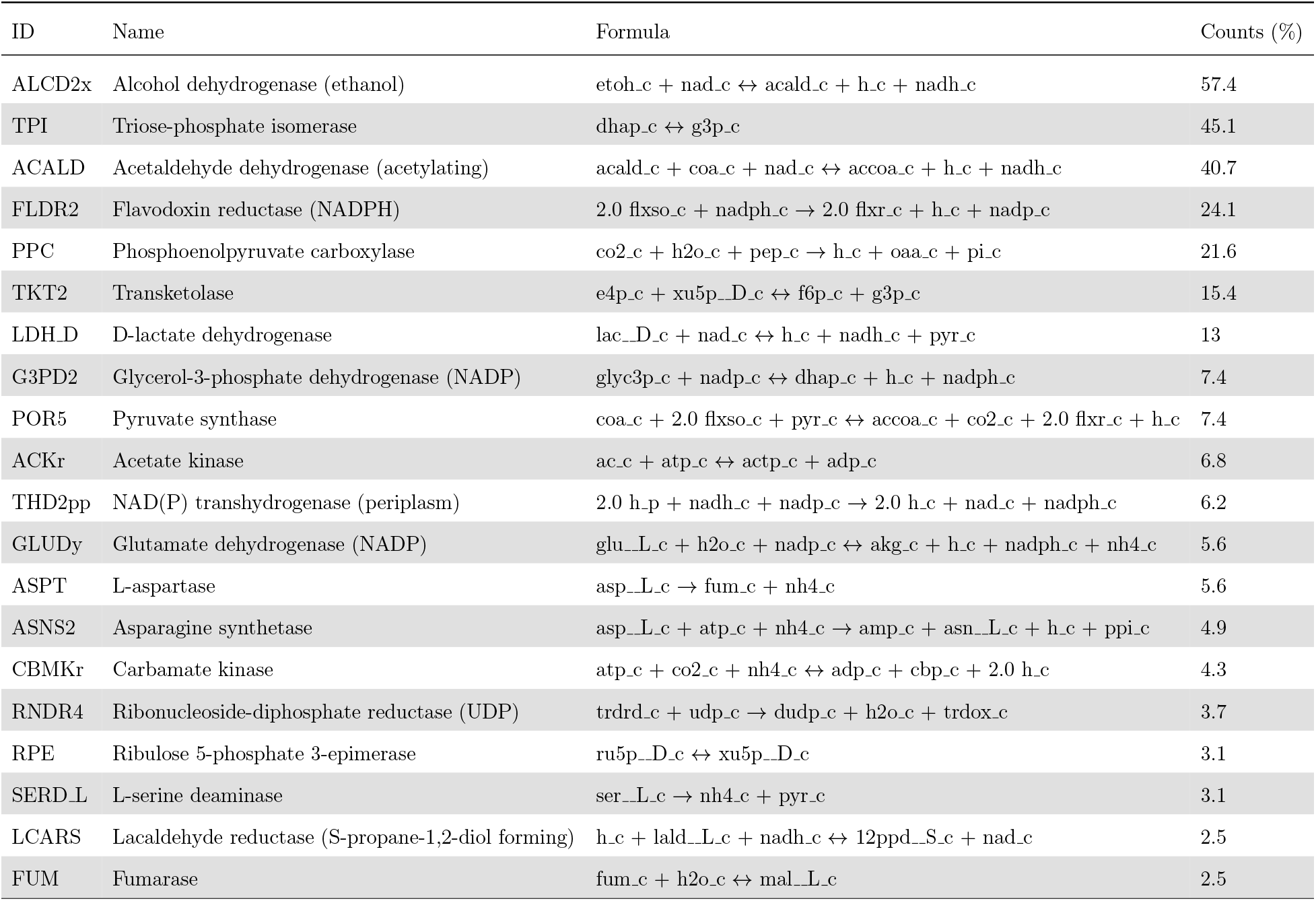
Top 20 reaction deletions for design parameters *α* = 5, *β* =1 with 162 designs. Counts indicate the percentage of designs where the deletion is used. All reaction and metabolite abbreviations used in this study correspond to BiGG identifiers (http://bigg.ucsd.edu).

##### Module reaction usage reveals pathway interfaces and unbiased module definition

The modular cell optimization formulation (Section 2.1) not only identifies genetic manipulations in the modular cell, but also in the production modules. Module reactions correspond to reactions deleted in the chassis but inserted back in specific production modules to enable compatibility. We examined the module reactions used by all designs (Figure 3). As expected, ALCD2x, ACKr, and LDH D, are used by ethanol, acetate, and lactate production networks, respectively. More notably, we also observed several reaction modules are used for specific products, for example, MDH and FUM 3-methyl-2-oxobutanotae and 2,3-dihydroxy-3-methylbutanoate, resepectively, that are naturally precursors of valine and artificially of isobutanol^37,38^. These module reactions likely play a role in both the synthesis of relevant TCA precursors and the secretion of succinate as an electron sink. Interestingly, fatty acids tend to use TPI, which as mentioned earlier, its deletion activates the methylglyoxal bypass lowering the overall ATP yield. The first step in fatty acid biosynthesis, acetyl-CoA carboxylase, requires one ATP per mol of malonyl-CoA, explaining the usage of TPI as a module reaction for this family of products. Overall, module reactions enhance the compatibility of a modular cell, leading to more efficient strategies and revealing potential metabolic flux bottlenecks that are not always directly upstream of the target product.

**Figure 3:**
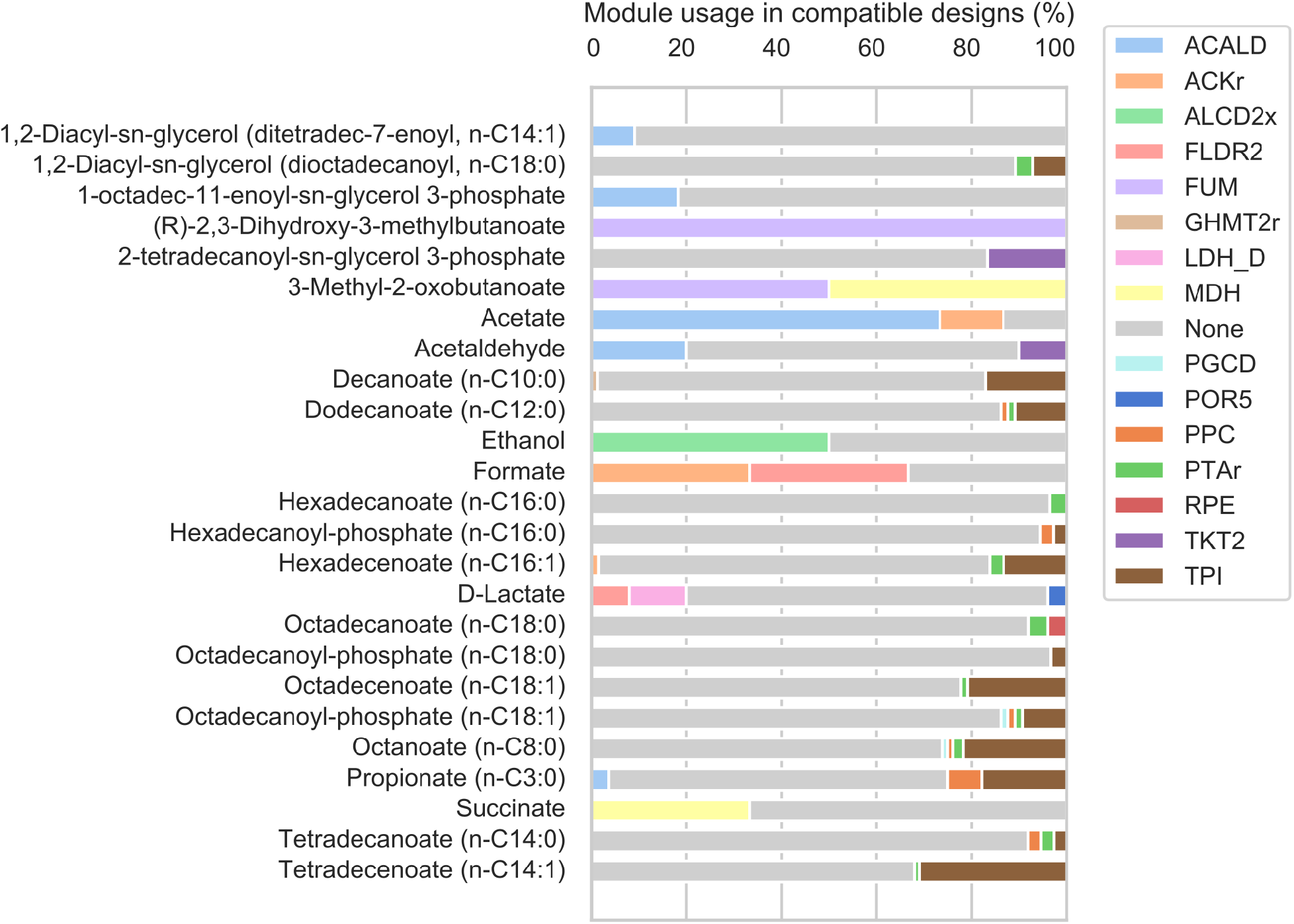
Module reaction usage for design parameters *α* = 5, *β* = 1. Only designs compatible with the product are considered in the module usage frequency.

##### Three modular cells is the smallest set needed to cover all compatible products

We next aimed to identify the smallest set of modular cells that include all compatible products in the library (Section 2.6.2). For the Pareto set of designs *α* = 5, *β* = 1, we enumerated a total of 12 minimal covers of size 3. These covers are spanned by combinations of 8 unique designs (Figure S5). We selected the cover k that contains designs 82, 121, and 124, which use few deletions and have similar genetic manipulations among them. All designs in this cover have in common the deletion of ALCD2x and LDH_D, disabling production of ethanol and lactate, the major reduced products of anaerobic growth in *E. coli*. Designs 121 and 124 have 57 compatible products in common, while design 121 is uniquely compatible with ethanol, formate, and 2,3-dihydroxymethylbutanoate, and design 124 is uniquely compatible with succinate (Figure 4 a). These two designs only differ in that design 121 uses FUM deletion while design 124 uses MDH deletion (Figure 4 b). Different from designs 121 and 124, design 82 is the only design that features the deletion of FLDR2 and PPC and is uniquely compatible with 24 modules, all for fatty acids synthesis. FLDR2 is coupled with POR5 to form a pathway for the reduction of pyruvate into acetyl-CoA consuming *nadph* (Figure 4 c), a key redox cofactor in fatty acid biosynthesis. PPC deletion is a metabolic engineering strategy to increase *nadph* available that has been experimentally validated.^31^ Overall, these designs can be efficiently built due to their similarity, and are mainly composed of strategies that have been demonstrated in isolation and cover large product families.

**Figure 4:**
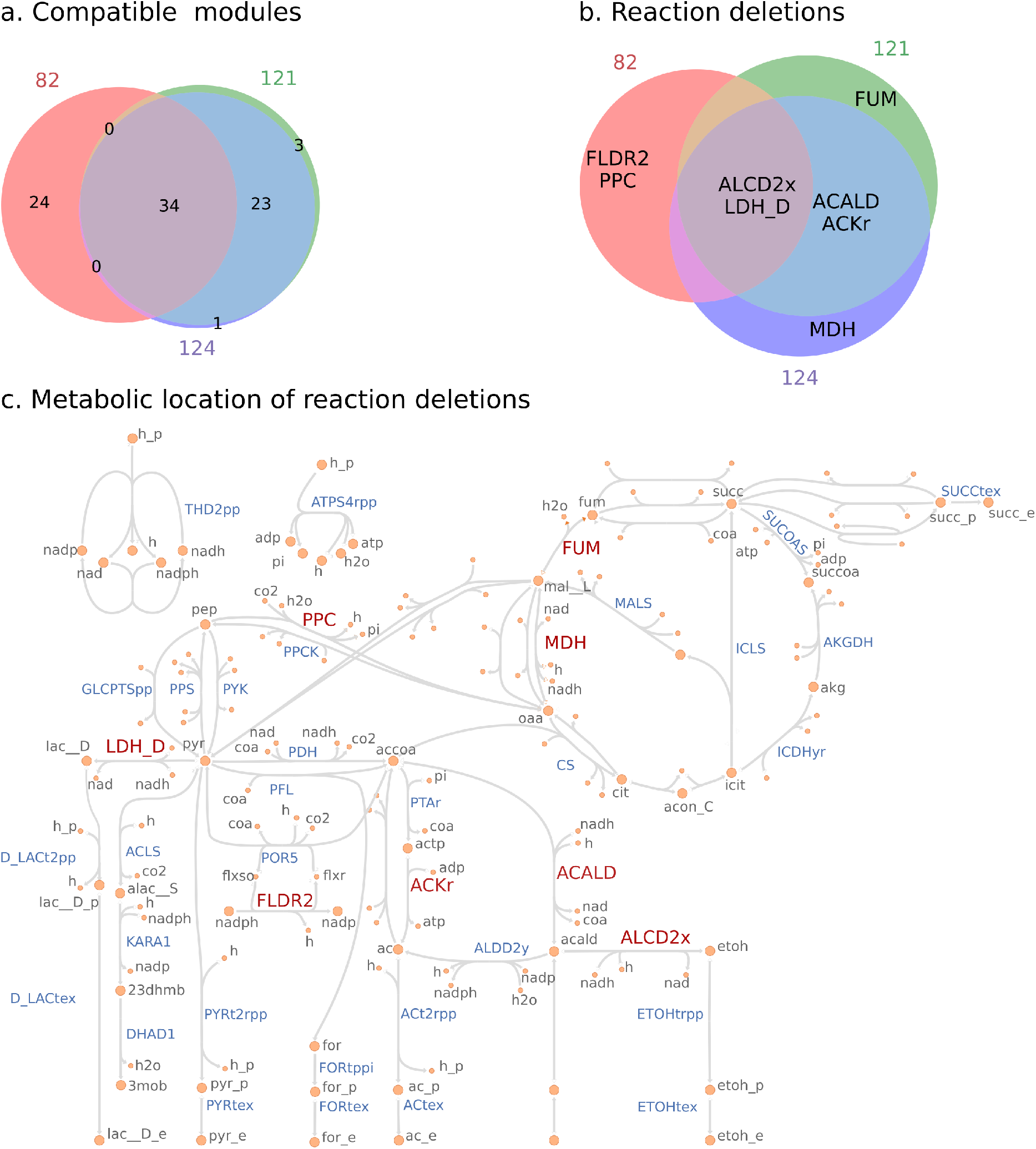
Comparison of the designs in the selected minimal cover. (a) Venn diagram of products compatible with each design. The products uniquely compatible with specific designs are (see http://bigg.ucsd.edu for abbreviation descriptions): Design 121: *etoh, for, 23dhmb*; Design 124: *succ*; Design 82: *pg140, 2hdecg3p, 2odec11eg3p, 1agpg180, pe140, pg161, pg141, 2hdec9eg3p, pgp161, 2agpg180, 1ddecg3p, pg120, pgp141, pgp140, pe141, ps140, apg120, ps120, pgp120, pe120, lipidX, 2tdecg3p, 2odecg3p, ps141*. (b) Venn diagram of reaction deletions that constitute each design. (c) Metabolic map with reaction deletions colored in red.

#### 3.3 Design of E. *coli* modular cells for conversion of hexoses and pentoses

##### Non-glucose carbon sources require more genetic manipulations for high compatibility designs

We designed modular cells to consume other relevant fermentable sugars besides glucose also present in biomass feedstocks, including pentoses (i.e., xylose and arabinose) and hexoses (i.e., galactose and mannose) (Figure 5 a). For this case study, everything remained the same except for the substrate uptake reaction in the model which was changed to reflect the sole carbon source in each case. We first scanned the distribution of design compatibilities resulting from various combinations of *α* and *β* for each carbon source (Figure S4 b-e). All cases plateaued at maximum compatibilities around 50%; however, galactose, arabinose and xylose required at least *α* = 10*, β* = 2 to reach this level, while glucose and mannose reached it with only *α* = 5*, β* = 1. Hence, we selected *α* = 10*, β* = 2 for further analysis. Overall, this simulation reveals the possibility of highly compatible modular cells for various hexose and pentose carbon sources, at the expense of an increased number of genetic manipulations for some of the carbon sources.

**Figure 5:**
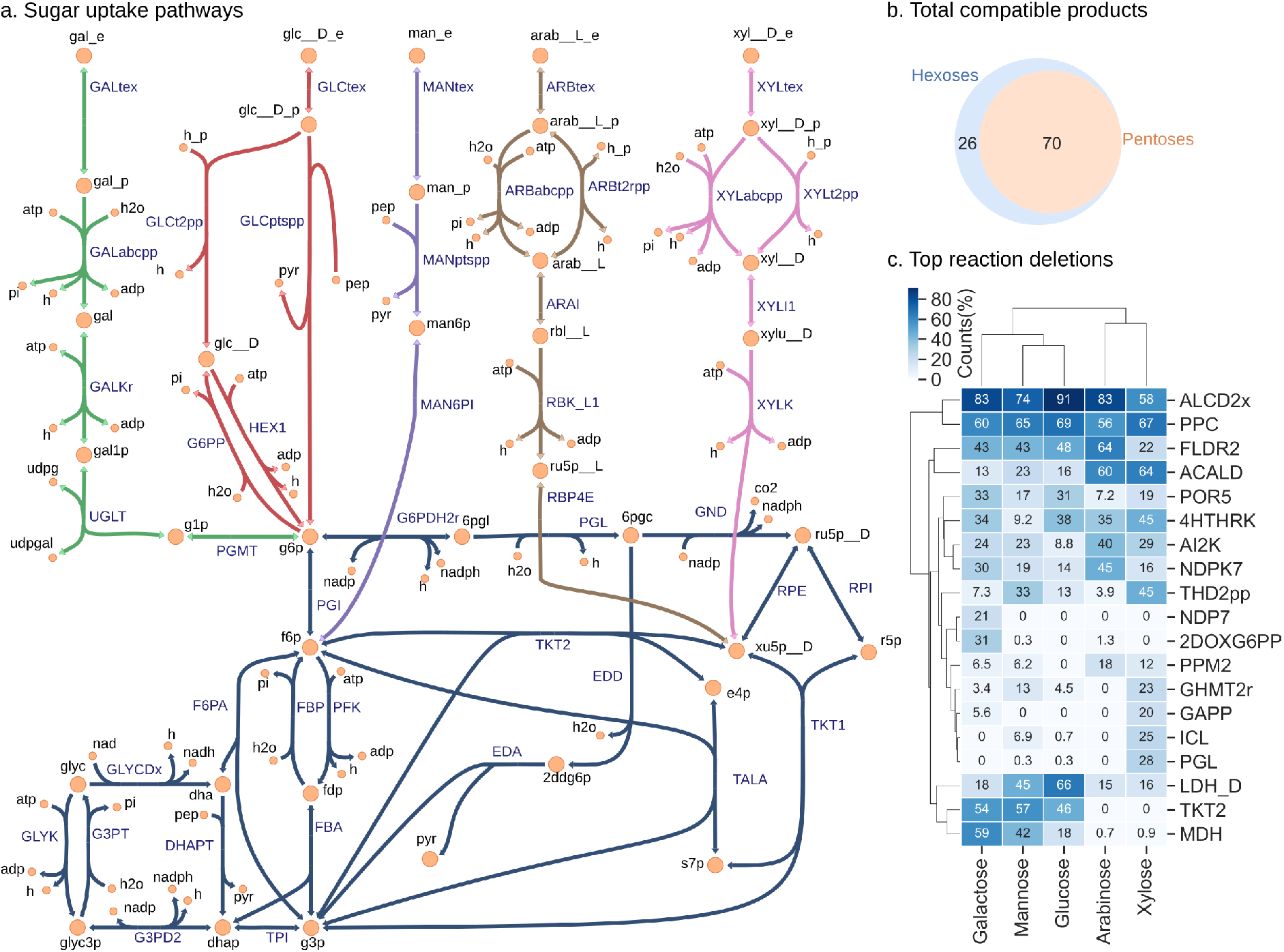
Design of modular cells for different carbon sources with design parameters *α* = 10, *β* = 2. (a) Sugar uptake, pentose phosphate, Entner-Doudoroff, and upper glycolysis pathways. (b) Venn diagram of total products compatible with designs using pentoses and hexoses. The 26 products uniquely compatible with hexoses are: *1agpg180, 2tdecg3p, 2agpg181, 3c3hmp, 3mob, 2hdecg3p, pe141, ps120, 1agpg160, 2agpg160, 23dhmb, ps141, 1agpe180, 2agpg180, apg120, 2agpe180, pe120, 2odec11eg3p, 4mop, lipidX, 3c2hmp, 2ippm, 2hdec9eg3p, 1agpg181, dha, 2odecg3p*. (c) Top 20 reaction deletions according to deletion frequencies average across carbon sources. The counts for each carbon source correspond to the percentage of designs containing that reaction deletion.

##### The effect of pentose uptake in redox metabolism leads to lower compatibility than hexoses

For the set of designs in each carbon source, we examined the total compatible products, i.e., the number of unique products compatible in at least one design from the Pareto front. This analysis revealed a group of 26 products (27% of the total 96 compatible products and 16% of the original library size) that are only compatible in designs with hexose carbon sources (Figure 5 b). The incompatibility of these 26 products is likely due to the lower reduction potential and different uptake pathways of pentoses with respect to hexoses (Figure 5 a). More specifically, analysis of the most deleted reactions in each carbon source revealed several differences in deletions between pentoses and hexoses (Figure 5 c). Notably, pentoses do not use TKT2 and MDH reaction deletions, while hexoses make highly frequent use of them. TKT2 is a key component of incorporating pentoses into glycolysis, and hence cannot be deleted by pentose consuming designs. MDH has been observed to be up-regulated under anaerobic conditions when the sole carbon source is pyruvate, galactose, or xylose with respect to glucose.^39^ Hence, MDH could be an important source of *nadh* for substrates with less reduction potential. Alternatively, MDH could also be important for *nadph* generation as part of a pathway involving NADP-dependent malic enzyme (ME2) that converts malate to pyruvate and generates one mol of *nadph*. Overall, pentose uptake does not use the oxidative branch of the pentose phosphate pathway, the most important source of *nadph* in *E. coli,^40^* hence limiting the products that can be growth-coupled to these carbon sources. Further study of the reactions that limit pentose compatibility could enable strategies to overcome it in certain cases (e.g., generation of alternative sources of _*nadph*_^41,42^_)._

#### 3.4 Compatibility towards modules unknown at the time of chassis design

##### Highly compatible designs are better suited to be re-purposed towards unknown products

To rapidly explore the large space of potential production modules, existing strains could be re-purposed for production of molecules not considered as part of the original design. To examine this scenario, we randomly partitioned the product library into two evenly sized groups, and independently used each partition as input for ModCell-HPC. This was done in triplicates, each corresponding correspond to a different random product partition. Hence, in each replicate there is a group of known products at the time of design and a group of unknown products. For the designs produced by ModCell-HPC, we computed their objective value and then compatibility towards unknown products, which we refer to as *unknown compatibility* of a design, a useful metric to understand the potential to re-purpose a given design. In contrast, *known compatibility* is the compatibility towards known products at the time of design, simply referred to as compatibility in previous cases study. The analysis of unknown compatibility of a new production module with an existing modular cell design is similar to the concept of degree of coupling that was previously introduced in MODCELL based on a different computation framework.^3^ The total number of designs for each product group and the unknown compatibility distributions noticeably changed across replicates (Figure 6 a). This result reveals the important effect of known products in the resulting designs, which could be further explored to identify “representative products” that can capture the necessary metabolic phenotypes required for certain product families. Remarkably, there was a high correlation between known and unknown compatibility of a given design (Figure 6 b-d). Hence, highly compatible designs are better suited to be re-purposed towards unknown products.

**Figure 6:**
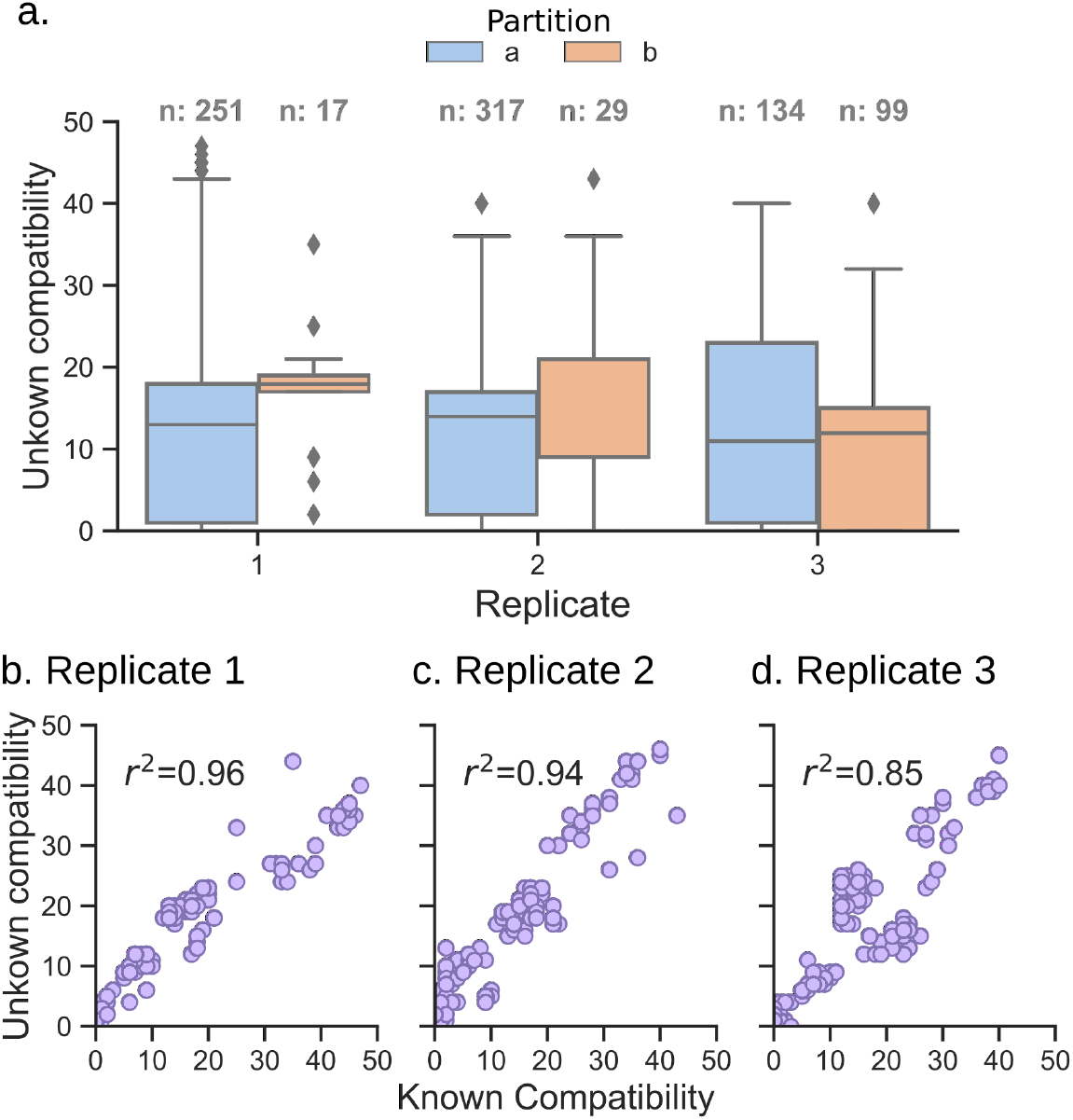
Compatibility towards unknown products in 3 random even partitions of the product library. (a) Distribution of unknown compatibility, n corresponds to the number of designs in each case. (b-d) Comparison between unknown and known compatibilities of each design for each replicate, where *r*^2^ is the Pearson correlation coefficient.

##### Deletion reactions that remove major fermentation byproducts and alter redox metabolism have the highest contribution towards unknown compatibility

To identify the specific genetic intervention strategies that contribute to the unknown compatibility of a design, we defined the unknown compatibility contribution of deletion reaction *j* (*ucc_j_*) as follows:

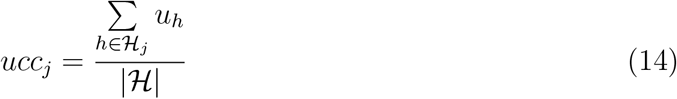

where 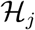 is the subset of designs from a ModCell-HPC solution (Pareto set 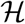) containing deletion reaction *j*, and *u_h_* is the unknown compatibility of design *h*. We computed *ucc* for all 3 replicates and examined the top 10 sorted by mean value (Table 3). The main contributors towards unknown compatibility were removal of major fermentative byproducts (lactate, ethanol, and acetate) followed by manipulation of redox pathways (THD2pp, FLDR2, MDH) and metabolic branch points (TKT2, PPC). Indeed byproduct removal strategies are the most common across the metabolic engineering literature.^28^ Strain re-purposing could be further explored with algorithms specialized for this task, e.g., by identifying module reactions in the unknown modules or using the existing strain as a starting point to identify genetic manipulations instead of a wild-type strain. In our analysis, we have identified that high modular cell compatibility and certain reaction deletions are positive indicators of compatibility towards unknown products.

**Table 3:** Top 10 reactions sorted by mean unknown compatibility contribution (*ucc*) among replicates (i.e., R.1, R.2, and R.3).

| ID | Name | $ucc$ | | | |
| --- | --- | --- | --- | --- | --- |
|  |  | R. 1 | R. 2 | R. 3 | Mean |
| LDH_D | D-lactate dehydrogenase | 13.2 | 10.5 | 11.9 | 11.9 |
| ALCD2x | Alcohol dehydrogenase (ethanol) | 11.5 | 10.5 | 11.8 | 11.3 |
| PTAr | Phosphotransacetylase | 4.0 | 4.8 | 6.5 | 5.1 |
| ACALD | Acetaldehyde dehydrogenase (acetylating) | 4.5 | 2.8 | 2.9 | 3.4 |
| THD2pp | NAD(P) transhydrogenase (periplasm) | 4.7 | 2.4 | 2.2 | 3.1 |
| ACKr | Acetate kinase | 3.8 | 2.2 | 1.7 | 2.6 |
| FLDR2 | Flavodoxin reductase (NADPH) | 2.0 | 2.2 | 2.9 | 2.4 |
| TKT2 | Transketolase | 2.6 | 2.0 | 2.5 | 2.4 |
| PPC | Phosphoenolpyruvate carboxylase | 2.3 | 2.2 | 2.5 | 2.3 |
| MDH | Malate dehydrogenase | 2.7 | 1.1 | 2.3 | 2.0 |

#### 4 Conclusions

In this study, we developed ModCell-HPC, a computational method to design modular cells compatible with hundredths of product synthesis modules. We applied ModCell-HPC to design *E. coli* modular cells with a product library of 161 endogenous metabolites. This resulted in many Pareto optimal designs for the production of these molecules, from which we identified three modular cells that include all compatible products. The designs feature strategies consistent with previous experimental studies aimed at optimizing production of a single product, reinforcing our confidence in the value of our simulations. Remarkably, the strategies not only include removal of major byproducts, but also modification of key metabolic branch-points. The modular cells were designed for growth-coupled production, which not only is expected to result in high product yields but also enables high-throughput pathway engineering approaches. Specifically, the modular cell can be simultaneously transformed with a module library to rapidly identify good candidates through adaptive laboratory evolution.^13,43^ We also used ModCell-HPC to design modular cells that utilize different hexoses and pentoses carbon sources. This revealed the limitations of pentoses towards coupling with certain products which might be addressed by redox cofactor engineering. Finally, we identified that high compatibility and certain reaction deletion are important features to re-purpose an existing modular cell towards new modules. Overall, ModCell-HPC is an effective tool towards more efficient and generalizable design of modular cells and platform strains that have recently captured the interest of metabolic engineers.^8^

## Acknowledgement

This research was supported by the NSF CAREER award (NSF1553250), the DOE BER Genomic Science Program (DE-SC0019412), and the Center of Bioenergy Innovation (CBI), U.S. Department of Energy Bioenergy Research Center supported by the Office of Biological and Environmental Research in the DOE Office of Science. The funders had no role in the study design, data collection and analysis, decision to publish, or preparation of the manuscript.

## Supplementary Materials

1. Supplementary Material 1 (SM1): Supplementary figures.

2. Supplementary Material 2 (SM2): Designs for selected parameters *α* = 5, *β* = 1.

