## Supplementary Material 1 (SM1) for "Computational Design and Analysis of Modular Cells for Large Libraries of Exchangeable Product Synthesis Modules"

### Supplementary Figures

Figure S1: Solution improvement process.

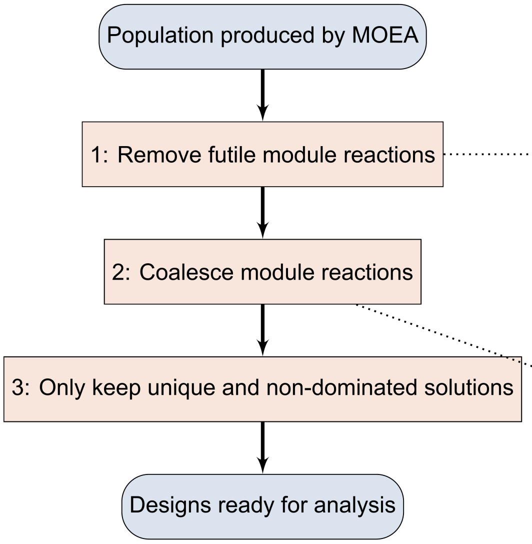


Futile modules reactions do not have an effect on the objective:

| Design | Deletions | M1 | M2 | O1 | O2 |
|---|---|---|---|---|---|
| Before removal | a,b,c | a,b | c | 0.8 | 0.5 |
| After removal | a,b,c | a | | 0.8 | 0.5 |

These are identified by systematically removing each module reaction for each objective, and if the objective value is diminished then the module reaction is kept and otherwise it is removed.

Consider the following designs with the same reaction deletions but different module reactions:

| Design | Deletions | M1 | M2 | O1 | O2 |
|---|---|---|---|---|---|
| 0 | a,b,c | a | | 0.5 | 0 |
| 1 | a,b,c | | b | 0 | 0.5 |

Both designs have the same deletions but independently found good module reactions, so the following design dominates both 0 and 1:

| Design | Deletions | M1 | M2 | O1 | O2 |
|---|---|---|---|---|---|
| 2 | a,b,c | a | b | 0.5 | 0.5 |

These are identified by condensing groups of designs with the same reaction deletions but different module reactions into the best possible design.

Figure S2: Chemical properties of the product library. DoR is the degree of reduction (mol $e^-$/ mol C), which is computed assuming a constant valency of 4, 1 , -2, and 5 for C, H, O, and P, respectively. For example, ethanol has 2 carbon atoms, a molecular weight of 46 g/mol, and a DoR of 6 (mol $e^-$/mol C). The molecular weight and the number of carbon atoms have a Pearson correlation coefficient (pcc) of 0.98, while DoR and the molecular weight only have a pcc of 0.42.

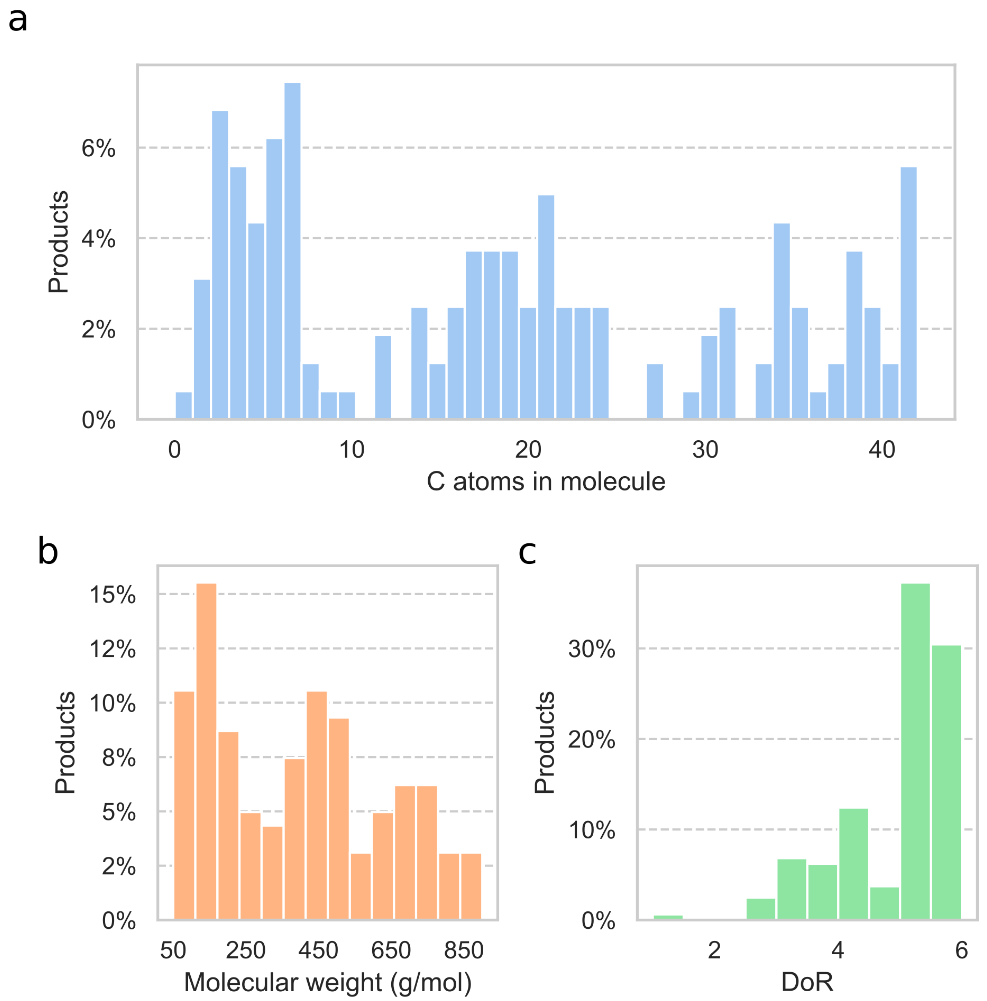

Figure S3: ModCell-HPC benchmarking with 20 products and design parameters $\alpha = 6$, $\beta = 1$. For a given run time, this analysis scans through all the combinations of migration interval, migration policy, and population size. Note that coverage values are not directly comparable between run times since they use a different reference Pareto front.

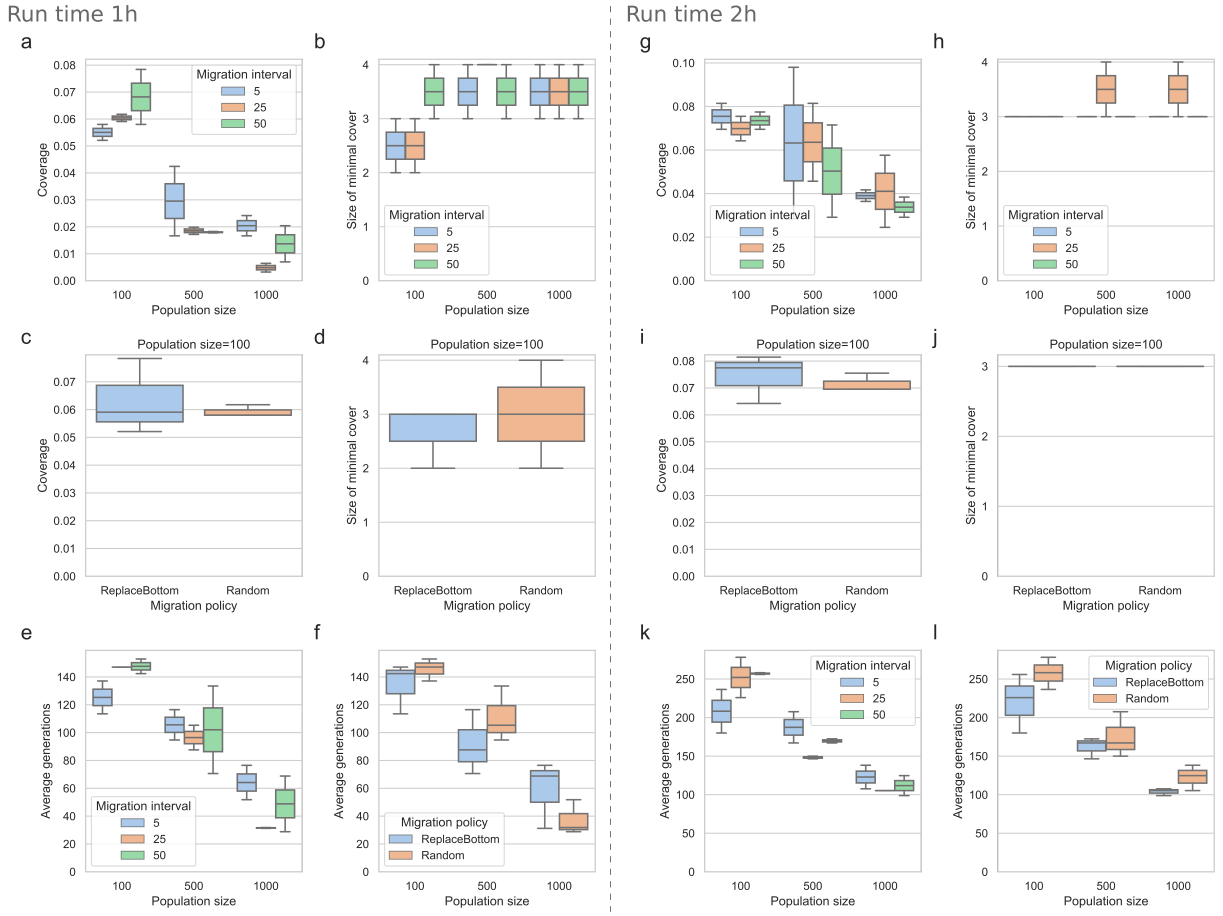

Figure S4: Compatibility of all designs in a Pareto front as a result of the design parameters. Each panel corresponds to a unique carbon source as the only difference in model configuration.

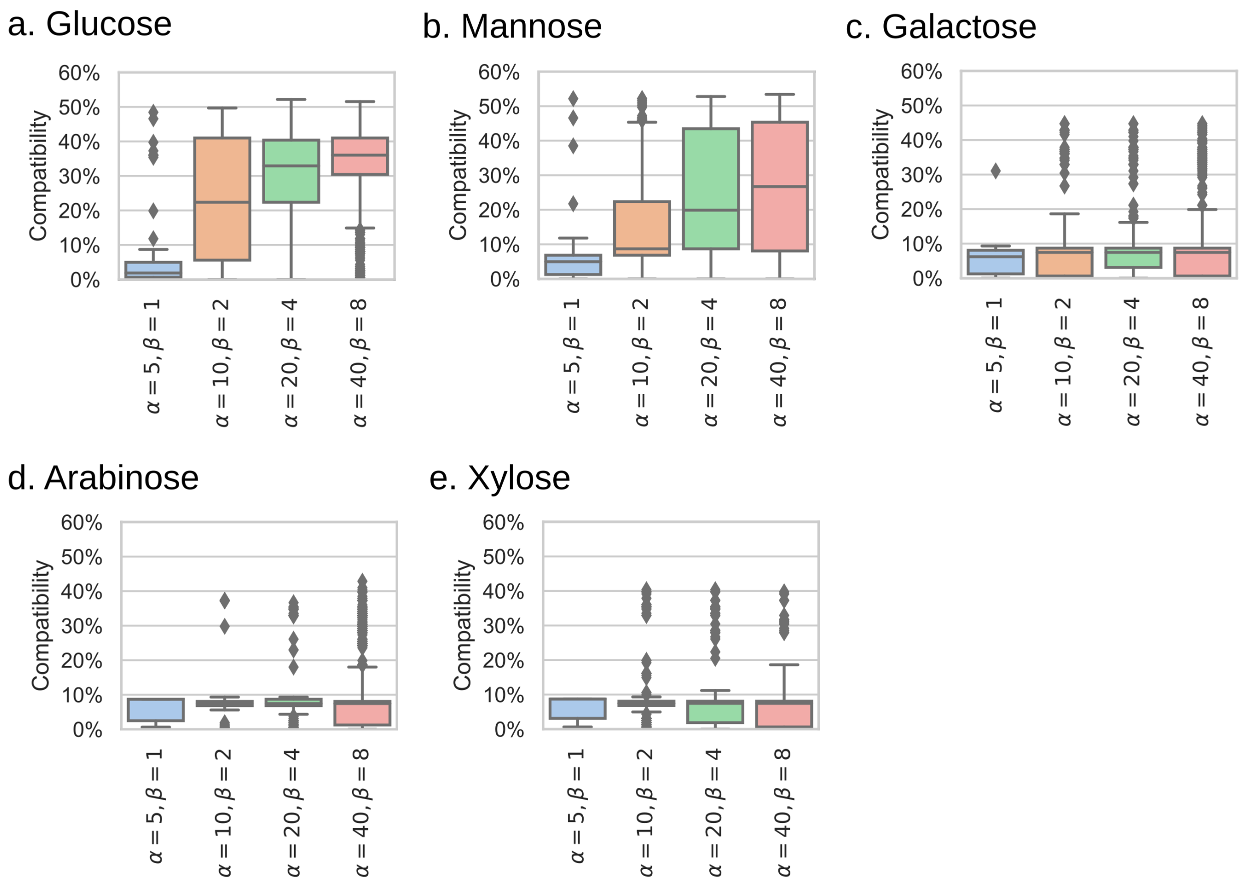

Figure S5: Bipartite graph representing minimal covers for design parameters $\alpha = 5$ and $\beta = 1$. Covers are colored in red and labeled with letters, while designs are colored in blue. All minimal covers are: a: [101, 109, 121], b: [109, 121, 124], c: [101, 110, 121], d: [25, 101, 121], e: [82, 101, 121], f: [110, 121, 124], g: [97, 109, 121], h: [97, 110, 121], i: [25, 97, 121], j: [25, 121, 124], k: [82, 121, 124], l: [82, 97, 121].

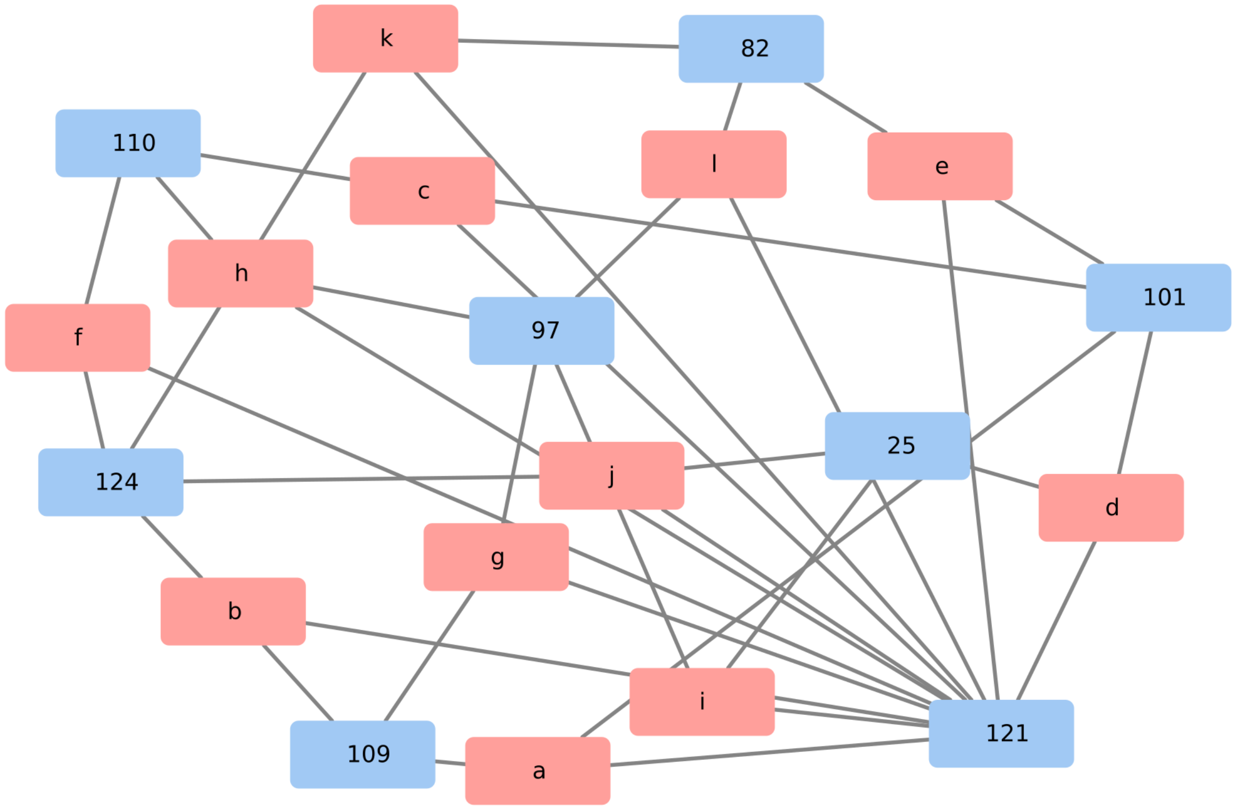
